# A Bayesian approach to mediation analysis predicts 206 causal target genes in Alzheimer’s disease

**DOI:** 10.1101/219428

**Authors:** Yongjin Park, Abhishek K Sarkar, Liang He, Jose Davila-Velderrain, Philip L De Jager, Manolis Kellis

## Abstract

Characterizing the intermediate phenotypes, such as gene expression, that mediate genetic effects on complex diseases is a fundamental problem in human genetics. Existing methods utilize genotypic data and summary statistics to identify putative disease genes, but cannot distinguish pleiotropy from causal mediation and are limited by overly strong assumptions about the data. To overcome these limitations, we develop Causal Multivariate Mediation within Extended Linkage disequilibrium (CaMMEL), a novel Bayesian inference framework to jointly model multiple mediated and unmediated effects relying only on summary statistics. We show in simulation that CaMMEL accurately distinguishes between mediating and pleiotropic genes unlike existing methods. We applied CaMMEL to Alzheimer’s disease (AD) and found 206 causal genes in sub-threshold loci (p < 10^−4^). We prioritized 21 genes which mediate at least 5% of local genetic variance, disrupting innate immune pathways in AD.

## Introduction

Alzheimer’s disease (AD) is a neurodegenerative disorder that evolves over decades and involves multiple molecular processes including accumulation of amyloid beta^1^, propagation of neurofibrillary tangles in the brain, and ultimately cognitive decline^2^. However, the precise gene-regulatory mechanisms and molecular pathways involved in AD progression remain to be elucidated. Case-control studies of differential gene expression in large postmortem brain cohorts have identified multiple genes and regulatory elements associated by AD^3–5^. The gene regulatory networks of these differentially-expressed genes have revealed promising associated genes for AD^3,4^, However, interpretation of case-control differential expression is challenging because it requires accounting for reverse causation of disease on gene expression, biological confounding, and technical confounding.

Large-scale meta analysis of genome-wide association studies (GWAS) for AD have identified 21 loci that influence AD susceptibility^6^. Unlike case-control differential expression, GWAS can in principle reveal causal relationships, because there is no possibility of reverse causation: late-onset complex disorders cannot change the genotypes at common genetic variants. However, the fundamental challenge in interpreting GWAS is that 90% of GWAS tag SNPs lie in noncoding regions^7^, making it difficult to identify the target genes and the causal mechanisms with relevant cell type information. Causal genes are not necessarily closest to the GWAS tag SNP, but may instead be distal to the causal genetic variant and linked by long-ranged chromatin interactions^8–10^.

Mendelian randomization^11,12^ resolves causal directions by using genetic variants as instrumental variables (IV) in causal inference. However, MR assumes no unmediated effecs on phenotype, no horizontal pleiotropy^11,13^ (the SNP affects both genes and phenotypes), and no LD-level linkage (two causal SNPs affecting genes and phenotypes in LD)^14^. A recent meta-analysis method, MR-Egger, partly relaxes these assumptions by modeling unmediated effects as a bias in regression of GWAS effect sizes on molecular QTL effect sizes^15^. However, it still assumes that genetic variants are not in LD (to perform meta-analysis) and that the estimated effect sizes reflect the true effect sizes of the variants (rather than being inflated by LD).

Recent methods for transcriptome-wide association studies (TWAS) have made important advances in identifying genes which could be causal. Unlike MR, TWAS aggregates information of multiple SNPs in LD to find genes whose cis-regulatory variants have correlated effect sizes for both gene expression and downstream phenotypes^16–18^. However, TWAS methods are fundamentally limited because they cannot distinguish between causal mediation, pleiotropy, linkage between causal variants, and reverse causation (Fig 1b), which could lead to inflated false positives as pleiotropy is quite prevalent in genetics^14^. Moreover, TWAS often finds multiple genes within a locus due to statistical correlations rather than independent biological effects because analysis is performed one gene at a time, ignoring gene-gene correlations^19^.

**Figure 1.**
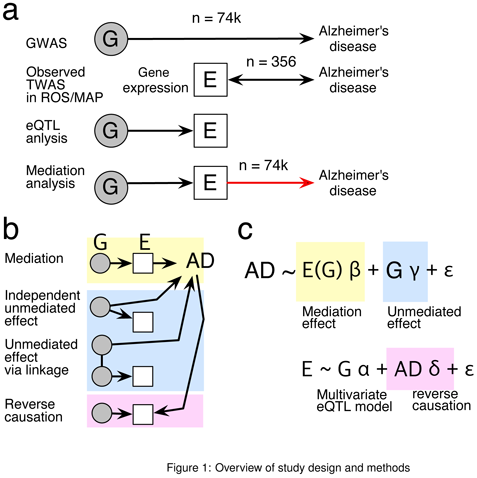
Schematic of the CaMMEL method. (**a**) Transcriptome-wide causal mediation analysis overcomes limitation of observed TWAS by taking advantages of large statistical power of Alzheimer’s disease GWAS statistics and brain tissue-specific regulatory contexts of eQTL data. (**b**) Correlation between gene expression and disease status can arise from mediation, pleiotropy and reverse causation. (**c**) CaMMEL jointly estimates two regression models taking into account multiple sources of gene expression variation.

Here, we present Causal Multivariate Mediation within Extended Linkage disequilibrium (CaMMEL), a new method for causal mediation analysis to find target genes from large-scale GWAS and molecular QTL trait association summary statistics (Fig 1a). With CaMMEL we address three aspects of the causal gene inference problem. First, CaMMEL leverages multiple SNPs in c/s-regulatory window to account for LD. Second, CaMMEL explicitly models mediation effect of multiple genes in close proximity to select a sparse set of causal genes that explains away non-causal correlated genes. Finally, CaMMEL models mediation effects while adjusting for unmediated effects. In simulation we found that CaMMEL correctly distinguishes mediating genes from pleiotropic genes, achieving higher statistical power than TWAS^17,18^ and Mendelian randomization with Egger regression (MR)^15^.

We applied CaMMEL to identify mediating genes for AD, combining GWAS of 74,046 individuals (25,580 cases and 48,466 controls)^6^ and brain gene expression of 356 individuals from the Religious Orders Study and the Memory and Aging Project (ROSMAP)^4,20^. We found 774 protein-coding genes with significant non-zero effect (FDR < 10^−4^), of which 206 are located in proximity of subthreshold GWAS SNPs (p < 10^−4^; Supp Tab 1), of which 66 explain at least 5% of local genetic variance. We further discussed 21 genes in subthreshold / GWAS loci that explain 5% of local variance, which includes the peroxisome regulator and cytochrome complex genes (*RHOBTB1*^21–23^, *CYP27C1* and *CYP39A1*) and several microglial genes (*LRRC23*^24,25^, *ELMO1*^26–28^, *RGS17*^29^, *CNTFR*^30^) which support functional roles of innate immune response in AD progression^2,31^.

## Results

### Overview of causal mediation analysis with multiple genes and SNPs

The goal of causal mediation analysis is to disentangle mediated (indirect) and unmediated (direct) effects from total effects between exposure and outcome variables^32^. Here, we define the mediated effect as causal transcriptomic regulatory mechanism of the available eQTL data, and broadly define the unmediated effects to include non-causal horizontal pleiotropy^11^, LD-level linkage^14^, and unidentified mediated effects of other regulatory mechanisms. Unlike conventional approaches, we assume phenotype data (AD) are generated with multiple mediators^33,34^ with coefficients *β* and the unmediated genetic effect with multivariate effect size *γ*; expression mediators are generated from multivariate eQTL effects *α* with unwanted reverse causation effect *δ* (Fig 1c).

A standard method is to perform regressions in two stages^33^: First, estimate the multivariate regression of the mediators (genes) on the genotypes adjusting for reverse causation and other types of mediator-exposure confounding; then estimate the multivariate regression of phenotype on imputed mediators (potential outcome) with putative unmediated genetic effects. We can independently correct for the confounding effects in gene expression matrix by half-sibling regression (Online methods)^35^, but the estimation of mediation effect sizes is fundamentally challenging because the mediator, e.g., gene expression, is usually not measured in the GWAS cohort, and imputation is not possible because the individual level genotypes required to impute the mediator are not available.

### Model specification of summary-based causal mediation analysis

CaMMEL explicitly models the probabilities of multivariate mediation effects and unmediated effects on traits by reformulating the multivariate mediation models into the equivalent summary statistic-based model based on Regression with Summary Statistics (RSS)^36^. The key idea of RSS is to reformulate a individual-level multivariate regression to a generative model of the observed univariate effect size vector 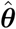 with the multivariate mean vector *θ* and the covariance matrix estimated from LD matrix *R* (Online methods; eq. 5). We developed an efficient approximate inference algorithm for the RSS model and validated that it performed competitively with regression models on individual-level genotype data such as Bayesian Sparse Linear Mixed Model^37^ (Supp Fig 3).

In CaMMEL, we model the total multivariate effect size vector *θ* by decomposing it into a linear combination of multivariate eQTL effect sizes *α_k_* for each gene *k* with the coefficients *β_k_*, and a multivariate effect size vector *γ* for the unmediated genetic effects:

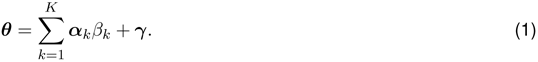

The true multivariate eQTL effects, *α_k_*, are *a priori* unknown, but can be inferred from the univariate eQTL effect sizes using RSS. In CaMMEL, we directly incorporate the RSS model for the univariate eQTL effect size vectors:

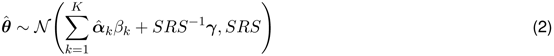

where *S* is a diagonal matrix of standard errors (Online methods). We assume the spike-and-slab prior on the vector of mediation effect sizes *β* in order to find a sparse set of non-zero effects from the correlated mediators^38–41^, but we assume a normal prior on direct effects *γ* to prevent the background unmediated effects from completely diminishing to zero.

### CaMMEL generalizes the MR and TWAS methods in theory

To gain mathematical insights into the CaMMEL model, we derived the coordinate-wise maximum likelihood estimate (MLE) of mediation coefficient on gene *k*, 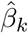 with estimated standard error 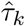 using the delta method^42^:

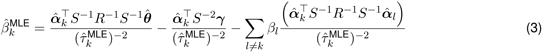

where estimated standard error 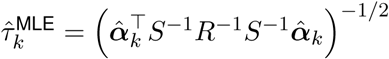. We describe technical details and the derivation in the supplementary texts.

With a large sample size and consistent minor allele frequency between GWAS and eQTL cohorts, we can interpret the first term as covariance between GWAS trait and gene *k*, the second term as covariance between unmediated effect and gene *k*, and the third term as summation of influence from all the other genes. Under the asymptotic normality, the Wald test statistic on *β_k_* converges to the standard normal distribution, 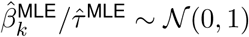.

TWAS^17^ is a special case of CaMMEL where there is no directional bias in unmediated effect, 𝔼[*γ*] = 0, and there there is no directional bias in mediated effects, 𝔼[*β_l_*] = 0 for all *l* ≠ *k*. MR^15^ is a special case of CaMMEL where there is no LD, *R* = *I*, no directional bias in mediated effects, but there could be directional bias of unmediated effects (which is modeled as a scalar parameter). As a consequence, TWAS and MR will only accurately estimate causal effect sizes if these conditions hold.

### CaMMEL performs well in simulations and correctly controls unmediated effect

We simulated gene expression data and Gaussian phenotypes using genotypes of 1,709 individuals on 66 real extended loci (Online methods). We compared the performance of CaMMEL with TWAS^18^ and the multi-SNP MR method^15^. Here, we show simulation results with two causal genes and two causal QTLs per gene as a representative example (Supp Fig 1), but we found similar trends with different number of causal genes and eQTLs per gene. When simulating with unmediated effects, we found CaMMEL achieved highest power and accuracy compared to TWAS and MR. At fixed FDR 1%, CaMMEL achieves 60% power to detect mediating genes when mediation explains 20% of genetic variance, and achieves nearly 50% power when mediation explains 10% of genetic variance (Supp Fig 1a). To investigate the robustness of methods against infused horizontal pleiotropy, we computed the percentage of falsely discovered genes at the threshold where each method achieved best precision and recall (maximum F1 score) in causal gene discovery (Supp Fig 1b). We found almost no evidence of spuriously associated genes in the discoveries made by CaMMEL, maintaining the fraction of false discoveries below 1%.

In contrast, TWAS, the statistical power and accuracy of TWAS degrade as we increase the level of unmediated effect. As expected by our analytical result, we found that TWAS had comparable power and AUPRC to CaMMEL only in the absence of unmediated effects (Supp Fig 1a). We confirmed weakened power of TWAS occurred when it confuses mediation with unmediated associated effect (Supp Fig 1b). We found unmediated associated genes are more frequently included in the top TWAS discoveries (5-10%) than CaMMEL (5%) and MR (2%). MR had essentially zero power in all of the simulated scenarios because the SNPs included in the model violate the independence assumption of the method (Supp Fig 1a).

### Transcriptome-wide mediation analysis in AD

We applied CaMMEL to AD GWAS and postmortem brain gene expression from the ROSMAP project to detect causal mediating genes in AD. We analyzed 2,077 independent, extended loci containing at least 100 well-imputed SNPs in the GWAS and ROSMAP. We allowed genes to span multiple LD blocks (based on SNPs within 1 Mb of the gene body) and empirically calibrated the null distribution of mediation by parametric bootstrap^43–46^. We controlled false discovery rate of the multiple hypothesis testing problem, estimating prior probability of alternative hypotheses from data^47^. Here, we defined cis-regulatory eQTLs and target genes with p-value < 0.05. We only considered genes with at least one eQTL after the p-value cutoff to avoid weak instrument bias in MR^48^. We included all GWAS SNPs in the background unmediated effect model if they appeared in the imputed genotype matrix, thus providing a conservative estimate of mediation effect, by allowing unmediated effect to explain away mediated effect.

We found 774 unique protein-coding genes with significant non-zero mediation effects (FDR < 10^−4^). Of them, we found 43 are located in genome-wide significant loci, 163 in subthreshold loci (p < 10^−4^; Fig 2a), and 568 in weakly associated regions (p > 10^−4^). Of the 43 genes in GWAS loci, in 9 cases we found the lead SNPs from GWAS agree with cis-eQTL effects, indicating the primary genetic effects are found in the postmortem brain tissues. In the remaining cases strongest effects may exist in different tissues and may only become visible in single cell eQTL data.

**Figure 2.**
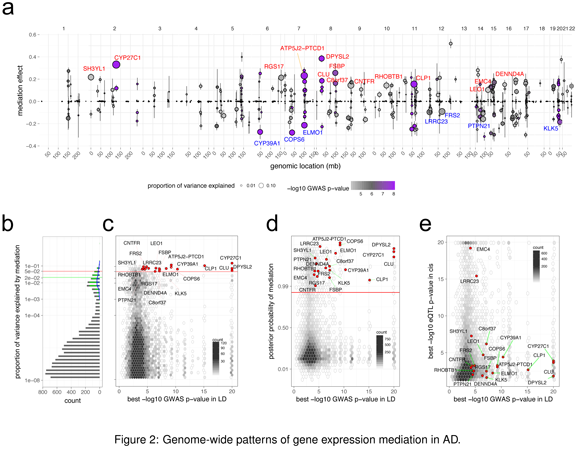
Transcriptome-wide mediation analysis reveals causal target genes in AD. (a) Mediation effect sizes over all the genes located in GWAS subthreshold region (p < 10^−4^). Size of dots are proportional to the proportion of local genetic variance explained by mediation. Errorbars show 95% confidence intervals. Dots are colored according to the GWAS significance. 21 genes (PVE > 5%) are annotated (Table 1). (b) Histogram showing proportion of variance explained (PVE) by gene expression mediation. *Gray bars:* total histogram of PVE; *green bars:* histogram of PVE for 774 significant mediation genes (non-zero mediation FDR < 10^−4^); *blue l/ne:* histogram of PVE for 206 significant mediation genes found in GWAS subthreshold regions (FDR < 10^−4^, GWAS p < 10^−4^). *Green horizontal line:* average PVE for 774 genes. *Red horizontal l/ne:* 5% cutoff for strong mediators. (c) Density estimation showing relationship between PVE (y-axis) and GWAS significance level (x-axis). The 21 strong mediators are shown as red dots. *Red horizontal line:* 5% cutoff. (d) Density estimation showing relationship between posterior probability of mediation (y-axis) and GWAS significance level (x-axis). We marked 21 strong genes with red dots. *Red horizontal i/ne:* posterior probability corresponding to FDR < 10^−4^ cutoff. (e) Density estimation showing relationship between best eQTL (y-axis) and best GWAS significance level (x-axis) within LD.

**Tables**

**Table 1.**
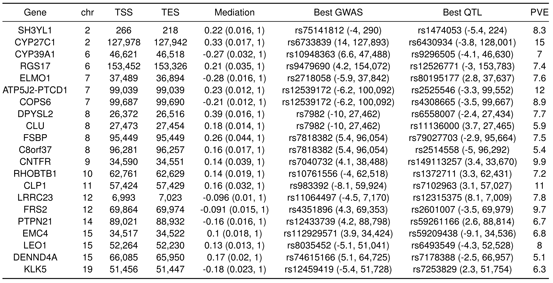
CaMMEL identified 21 genes explains at least 5% of local variance in subthreshold / GWAS regions. All genes in the list passed stringent p-value threshold < 3e-06 (FDR < .007%) and GWAS regions contain at least one subthreshold or more significant SNP (p-value < 10^−4^). *Gene:* gene symbol in `hg19`; *chr*: chromosome name; *TSS:* transcription start site with respect to strand (kb); *TES:* transcription end site (kb); *Mediation:* gene expression mediation effect size with standard deviation and posterior inclusion probability (first and second numbers in the bracket). *Best GWAS:* best GWAS ID in the LD block with z-score and location in kb. *Best QTL:* best eQTL SNP ID with z-score and location in kb. *PVE*: proportion of local genetic variance explained by gene expression mediation (in %).

We found on average the significant 774 genes each explain more than 2% of local genetically-driven phenotypic variance (proportion of variance explained, PVE). This proportion was similar for the 206 genes in the subthreshold/GWAS loci. However, the remaining genes explain essentially zero variance (Fig 2b). We found no clear evidence of correlation between GWAS significance with the proportion of local variance explained by mediation effects (Fig 2c) or the posterior probability of non-zero mediation effects (Fig 2d). Our results can be interpreted through omnigenic perspective^49^ that a large number of causal genes accounting for weak polygenic effects do not yet reach genome-wide significance with current cohort size, suggesting causal variants of mediation genes located in the subthreshold regions can eventually reach genome-wide significance level as we increase sample size.

### CaMMEL recapitulates known Alzheimer’s disease genes

The 774 identified mediating genes include several known AD genes that are significant mediators and co-localized in genome-wide significant loci (p < 5e-8). *COPS6* accounts for nearly 9% of the variance (Fig 3a) and the overall eQTL effect sizes are negatively correlated with AD susceptibility (-0.21 ± 0.012; Tab 1). *COPS6* constitutes the *COP9* signalosome (CSN) complex, and the biological function of CSN was explored regarding innate immunity. The CSN complex is evolutionary conserved, and plays a key role in plant innate immunity^50^. Recently conditional knock-out experiments on the other subunit *COPS5* conducted in macrophages show highly correlated activities with anti-inflammatory pathways^51^.

**Figure 3.**
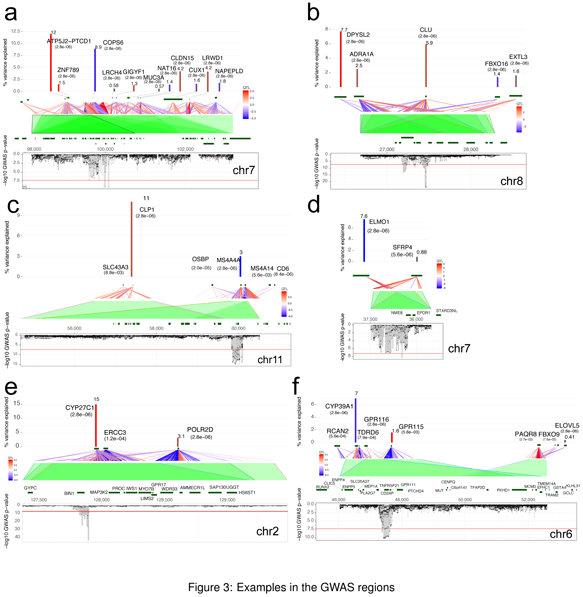
Local views of significant gene expression mediation on GWAS significant regions. In each plot, proportion of gene expression mediation (%) and GWAS Manhattan plot (upside down) are aligned according to genomic locations. eQTL links are colored based on z-score of effect sizes (blue = negative; red = positive). Ranges of correlation due to LD were covered with green shades. Gene bodies are marked by dark green bars in genomic location. Numbers within brackets denote mediation test p-value.

*CLU* explains nearly 6% of the local variance in the genome-wide significant locus (Fig 3b), and its mediation effects are positively correlated with AD susceptibility (mediation coefficient 0.18 ± 0.014; Tab 1). It is shown that *CLU* (clustrin) expression is elevated in AD brain and directly interacts with amyloid beta plaques, and interferes with clearance of neuritic plaques^52–55^.

*MS4A4A* explains 3% of the local variance with negative correlation with AD risk (Fig 3c). Although the functional role of *MS4A* complex is largely unknown^52,53^, recently *MS4A4A* was characterized as a novel surface marker for immature dendritic cells^56^.

### CaMMEL finds new genes in AD GWAS loci

We found 21 genes that explain high local genetic variance (at least 5%) in the GWAS / subthreshold regions (Fig 2a and Tab 1), of which 8 are located in the GWAS regions. A read-through gene *ATP5J2-PTCD1* mediates 12% of the local variance. The gene body is displaced from the strong GWAS peak in chromosome 7, but *cis*-eQTLs are located within the peak and share strong correlation within the LD (Fig 3a). Function of the readthrough protein is not well-understood, but *PTCD1* modulates mitochondrial precursor RNA and tRNA^57^ and the related protein *PTCD2* was already recognized as a biomarker due to its elevated expressions in AD^58,59^.

*DPYSL2* is distal from the strong GWAS peak in chromosome 8, but its *cis*-eQTLs show a large degree of overlap with the GWAS region in LD (Fig 3b). *DPYSL2* was up-regulated in cortex, striatum and hippocampus after ischemic stroke^60^, and it was listed among genes that modulate activation of microglial cells^61^. In fact, CaMMEL found that *DPYSL2* is positively correlated with AD progression.

*ELMO1* is not co-localized with the strong GWAS peak in chromosome 7, yet c/s-eQTLs are located within LD with GWAS SNPs (Fig 3d). This gene confers risk of immune disorders such as multiple sclerosis in previous GWAS^27,28^, and triggers phagocytosis in microglia interacting with other proteins^26^. Our mediation analysis demonstrates that brain tissue-specific causal mechanism of *ELMO1* is negatively correlated with AD risk, and implicates mitigated activity of clearing neuritic plaques.

We note several cases where CaMMEL points to biologically important genes which are not captured by GWAS. For instance, *cis*-eQTLs of *CLP1* are located almost independent of the significant GWAS region on chromosome 11, but the gene’s effect explains 11% of local variance (Fig 3c). *CLP1* is an interesting gene to follow up validation for neurodegenerative disorders; its mutation leads to damages in peripheral and central nervous systems altering tRNA synthesis^62,63^, which conveys consistent implication as *ATP5J2-PTCD1.*

Two cytochrome P450 complex genes^64^, *CYP27C1* (chr2) and *CYP39A1* (chr6) explain 15% and 7% of the local variance. *CYP27C1* is closely located with well-known *BIN1*, but the gene body does not overlap with strong GWAS signals. Instead, *cis*-eQTLs within 1Mb window cover that region and extend through non-zero LD effect of *CYP27C1* to nearly 2Mb around the GWAS locus (Fig 3e). The contribution of *BIN1* to AD is somewhat controversial^65–67^, and we expect follow-up mediation analysis with cell type-specific eQTLs will further clarify the mechanisms.

The gene body of *CYP39A1* does not overlap with GWAS SNPs; however, *cis*-eQTLs are in LD with the GWAS variants (Fig 3f). *CYP27C1* is involved in lipid metabolism^68^ and plays an important role in photoreceptors^69^. *CYP39A1* is involved in the generation of 25-hydroxy cholesterol from cholesterol. A regulatory role of hydroxy cholestrol in innate immunity is increasingly recognized by recent studies^70^, and a murine experiment showed that inflammation in central nervous system due to disrupted lipid metabolism down-regulated activities of cytochrome P450 complex^71^.

### CaMMEL finds strong mediators in subthreshold loci

Of the 21 strong mediators, 13 are located in the subthreshold regions. We highlight three examples in the subthreshold regions. *SH3YL1* is located at the tip of chromosome 2 containing small number of genes and gene body is enclosed within subthreshold GWAS region (Fig 4a). *SH3YL1* was found significant in GWAS with attempted suicide and expressed in brain, but underlying mechanisms are unknown^72^. Interestingly this gene was also found robustly associated height in previous TWAS^17,18^. Several previous researches noted subtle relationship between AD and suicide attempt^73,74^ and height^75^. However we think joint analysis with multiple GWAS traits will further edify detailed causal directions.

**Figure 4.**
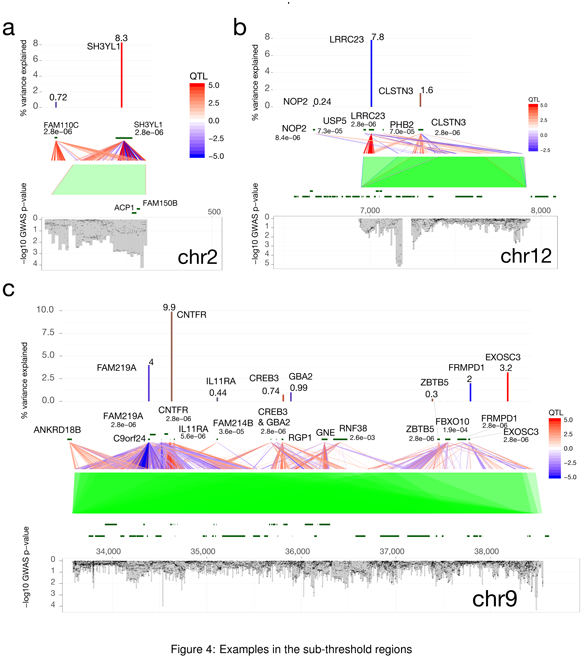
Local views of significant gene expression mediation on subthreshold regions. In each plot, proportion of gene expression mediation (%) and GWAS Manhattan plot (upside down) are aligned according to genomic locations. eQTL links are colored based on z-score of effect sizes (blue = negative; red = positive). Ranges of correlation due to LD were covered with green shades. Gene bodies are marked by dark green bars in genomic location. Numbers within brackets denote mediation test p-value.

*LRRC23* has *cis*-eQTLs in chromosome 12 between 7Mb and 8Mb that explain 7.8% of local genetic variance, followed by *CLSTN3* explains 1.6% (Fig 4b). Both mediation effects of *LRRC23* and *CLSTN3* are significantly nonzero. *LRRC23* and its *cis*-eQTLs were suggested as a modulator of innate immune network in mouse retina in response to optic neuronal injury^24^ and enriched in ramified microglial cells^76^. *CLSTN3* (calsyntenin-3) interacts with neurexin and forms synpatic adhesion^77^ and up-regulated by increase of amyloid beta protein and accelerated neuronal death^78^.

*CNTFR* explains 9.9% of genetic variance within 5Mb region (from 33Mb to 38Mb on chromosome 9) and multiple gene expression mediators are found (Fig 4c). Interestingly GWAS signals are also widely distributed with sporadic narrow peaks. *CNTFR*, interacting with *CNTF* and interferon gamma, stimulates murine microglia and increases expression of *CD40* on the cell surface^30^. Second best gene in the region, *EXOSC3* (RNA exsosome component 3), is also interesting. Exome sequencing followed by functional analysis in zebra fish identified that mutations in *EXOSC3* is causal to hypoplasia and spinal motor neuron degeneration^79^, and expression of *EXOSC3* was differentially regulated in human monocyte and macrophage induced by lipopolysaccharide, suggesting functional roles in inflammatory bowel disease^80^.

## Discussion

We developed novel Bayesian machine learning approach to causal mediation analysis that infers regulatory mechanisms from summary statistics of GWAS and eQTLs. Our approach borrows great ideas from two existing statistical approaches, MR-Egger regression and TWAS. MR-Egger adjust for unmediated effects by meta-analysis of effect sizes^15^, and TWAS correctly aggregate multiple SNP information taking into accounts of LD. However, these methods only provide partial solution of causal gene discoveries, posing strict assumptions such as independence between SNPs (MR-Egger) or giving up identifiability between mediated and unmediated effects (TWAS). On the contrary, CaMMEL addresses overall aspects of the causal mediation analysis in tight LD structure^36^, by explicitly modeling both mediated and unmediated effects of multiple genes in a unified Bayesian framework and to identify causal pathways by carrying out Bayesian variable selection^38^ between potentially collinear mediators^40,41^. Our direct Bayesian approach poses computational challenges on posterior probability calculation, but our efficient implementation made the method generally applicable to other related summary-based mediation and regression analysis.

Previous MR analysis on DNA methylation^81^ used stringent FWER cutoff on instrument variable selection to avoid weak instrument bias observed in MR on individual-level data^48^. Here we retained *cis*-regulatory eQTLs and target genes with lenient p-value cutoff (0.05) for several reasons. Since CaMMEL estimates the RSS model and it reduces the risk of double counting evidences from multiple weak IVs. Forever, unlike endophenotypes in epidemiological studies such as Body Mass Index, strong genetic association of molecular QTL include effects from neighboring SNPs, and therefore stringent p-value cutoff does not make the QTL a strong iv^27,82,83^. In fact, our approach to analyze multiple mediators make causal mediators tested in more strict settings when we include as many competitor genes as possible.

We established top priority list of 21 genes as strong causal mediators of AD. Our results suggest contribution of innate immunity in AD progression pointing to a specific set of genes with proper brain tissue contexts. We expect our analysis framework can provide more complete pictures by combining with cell-type and tissue specific eQTL summary statistics. Currently we assume mediators (genes) act independently conditioned on genotype information, but extensions of LD blocks to pathways and multiple tissues would need more parsimonious models, for instance, by sharing information across tissues and genes of through factored regression models^84^.

## Online methods

### Review of regression with summary statistics

A summary association statistic refers to a univariate effect size (a regression slope of a simple regression or logodds ratio in case-control studies) measured on each SNP without taking into account of LD structure. On a single trait GWAS, we normally have a vector of *p* summary statistics, effect size 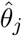 and corresponding variance 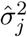 for each SNP *j* ∈ [*p*]^85^. However, due to LD (correlations between neighboring SNPs), an effect size measured on each single variant contains contributions from the neighboring SNPs. Unless we estimate causal variants by fine-mapping, inferring the multivariate effect from the univariate one, the univariate effect size of a SNP should not be interpreted as unbiased estimation of SNP-level signals^85,86^.

Here our interest is on the multivariate (true) effect size vector *θ* across all SNPs in the locus of interest. In principle, fixed effects can be modeled by a multivariate regression model, and parameters of this large regression model correspond to the multivariate effect *θ*. We assume an *n*-vector of individual-level phenotypes y was generated from *n* x *p* genotype matrix *G* on *p* SNPs with the multivariate effect sizes *θ*, and isotropic Gaussian noise parameter *σ*^2^. More precisely,

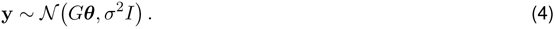

The regression with summary statistics (RSS) model^87,88^ provides a principled way to describe a generative model of the GWAS summary data. We assume the observed GWAS summary effect sizes 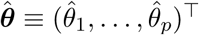 were generated from the same multivariate effects *θ* of individual-level multivariate GWAS model (eq. 4).

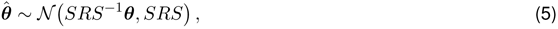

where we denote the *p*×*p* linkage disequilibrium (LD) matrix *R* and the diagonal matrix of expected squares, *S* with each element 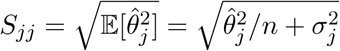. In practice we estimate the LD matrix 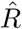 from a standardized genotype matrix of reference cohort by taking crossproduct, 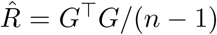. While other methods regularize 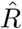 matrix by adding ridge regression penalty on the diagonal matrix^89^, here we regularized 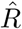 by dropping variance components corresponding to small eigenvalues (λ)^90^. We use a fixed cutoff λ < 10^−2^, but our results were robust to the different cutoff values such as λ < 10^−3^ and λ < 10^−1^.

### Derivation of the causal mediation model in RSS

We derive the mediation model (eq. 2), assuming that the observed univariate eQTL effects on a mediator *k*, 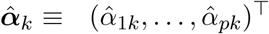 are generated from the RSS model with mean 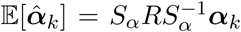 with expected squares *S_α_* of eQTL effects. Here we assume standard errors of GWAS effects can be substituted with standard errors of QTL effects up to some scaling factor, i.e., *S_α_* = *cS* with some real number *c* > 0, because association statistic is mainly determined by the underlying minor allele frequencies and samples sizes, and this allows us to rewrite the expectation of mediation effect (eq. 1):

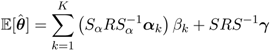

where the constant factor *c* cancels. With no measurement error, i.e., 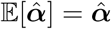, we can reasonably assume:

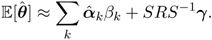

Substitution of this mean effect to the RSS model (eq. 5) yields the probabilistic model for CaMMEL (eq. 2).

### Variational Bayes inference of the mediation model

A key challenge in fitting the causal mediation model (eq. 2) is dealing with the covariance matrix in the likelihood. We exploit the spectral transformation of the LD matrix to make the model inference more amenable^91^. We take singular value decomposition of standardized genotype matrix, (*n*)^−1/2^*G* = *UDV*^T^, and decompose LD matrix into 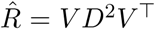, exposing that the effective number of sample size *ñ* is bounded by the sample size of reference panel, *ñ* < *n*, after the regularization 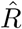. Defining 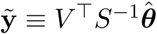 we obtain equivalent, but fully factorized, multivariate Gaussian model

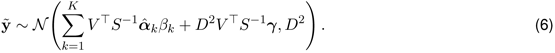

We efficiently fit the model using black box variational inference^92,93^ with a novel reparameterization trick^84,94^. Normally a high-dimensional multivariate regression is intractable problem since the total amount of model variance blows up as a function of dimensions (*p*). However, instead of dealing with large number of parameters (*p* SNPs), we deal with smaller *ñ*-dimensional aggregate random variables (*ñ* < *n* ≪ *p*), a linear combination of *p*-dimensional effects. More precisely we define

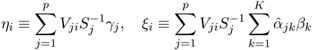

to rewrite the transformed log-likelihood of the model for each eigen component *i*:

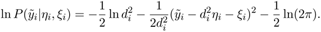

This reformulation not only achieves faster convergence by reducing variance^94,95^, but we also gain computational efficiency that we can sample all the eigen components independently in parallel taking full accounts of the underlying LD structure between SNPs.

The overall algorithm proceeds as follows: We first update surrogate distributions of *q*(*ξ*) ≈ 𝒩(*μ_ξ_*, *v_ξ_*) and *q*(*η*) ≈ 𝒩(*μ_η_*, *v_η_*) by minimizing Kullback-Leibler (KL) divergence between the surrogate *q* and true distribution *P*, *D*_KL_(*q*||*P*), by taking stochastic gradient steps with respect to the mean ∇_*μξ*_, ∇_*μη*_ and variance ∇_*vη*_, ∇_*vη*_^93^; we then then back-propagate this gradient to the gradient with respect to the original mean ∇_*μβ*_, ∇_*μγ*_ and variance parameters ∇_*vβ*_, ∇_*vγ*_ to eventually find *q*(*β*) ≈ 𝒩(*μ_β_*, *v_β_*) and *q*(*γ*) ≈ 𝒩(*μ_γ_*,*v_γ_*). We formulate the variational mean *μ_β_* and variance *v_β_* of the spike-slab distribution following the previous derivations^39^ (see our technical paper^84^ for details).

### Calculation of proportion of variance explained

We measured explained variance according to the definition of Shi *et al.*^90^ Total variance can be decomposed into mediated component, (*αβ*)^T^ *R*(*αβ*) and unmediated component (*γ*)^T^ *R*(*γ*). We checked this provides a reasonably tight lower-bound of actual model variance through simulations. Parameters, *α,β, γ*, are estimated by posterior mean of variational Bayes inference.

### Parametric bootstrap

Variational inference may only capture one mode of the true posterior, and so tends to underestimate posterior variances. Therefore, we do not rely on the Bayesian posterior variance to assess confidence in the estimated mediated and unmediated effect sizes. Moreover, the effects tend to correlate with each other then can lead to a severe collinearity problem. Instead, we use the parametric bootstrap to estimate the null distribution of these parameters and calibrate false discovery rates^43,44,46^.

### Confounder correction in gene expression matrix

We learn and adjust for unobserved non-genetic confounders in gene expression by half-sibling regression (Supp Fig 2)^35^. For each gene we find 105 control genes, selecting the most correlated 5 genes from each other chromosome. We assume cis-regulatory effects only occur within each chromosome and that inter-chromosomal correlations are due only to shared non-genetic confounders. This assumption is invalid when *cis*-regulatory variants under consideration have *trans*-effects on the control genes. Therefore, we regress out genetic effects on the established control genes. Half-sibling regression simply takes residuals after regressing out the control gene effects on observed gene expressions.

### Establishing the extended LD blocks

We establish independent LD blocks of SNPs within a chromosome and restrict mediation analysis within each block. We computed pairwise correlations between pairs of variants in the Thousand Genomes European samples within 1 megabase and with *r*^2^ > 0.1. We pruned to a desired threshold by iteratively picking the top-scoring AD GWAS variant (breaking ties arbitrarily) and removing the tagged variants until no variants remained. At convergence each small block is tagged by its top-scoring variant, and merge LD blocks overlapping in genome.

### Simulation framework

We used 66 real extended loci, randomly sampling 3 per chromosome that each harbored more than 1,000 SNPs. Within each region, we assigned each non-overlapping window of 20 SNPs to a synthetic gene. For each gene, we varied the number of causal *n_q_* SNPs (per gene) and generated *n_g_* gene expression data from a linear model with proportion of variance mediated through expression (PVE) from *τ* = 0.01 to 0.2. For each region, we selected *n_d_* SNPs to have a direct (unmediated) effect on the phenotype and selected genes to have a causal effect on the phenotype, varying the variance explained by the direct effect *δ* (horizontal pleiotropy^13^). For simplicity we used *n_d_* = *n_q_*, meaning direct effect contributes variance of one gene, and gene expression heritability was fixed to *ρ* = 0.17 as used in our previous work^96^. We carefully selected pleiotropic SNPs from the eQTLs of non-mediating genes. We evaluated the statistical power of each method at fixed FDR 1% and the precision by the area under the precision recall curve (AUPRC).

We used a generative scheme proposed by Mancuso *et al.*^18^ with a slight modification that includes unmediated genetic effects on the trait. Let *n_q_* be number of causal SNPs per each genetic gene; *n_g_* be number of causal mediation genes; *n_d_* be number of SNPs driving unmediated effect on trait. For each simulation, three PVE (a proportion of variance explained) parameters are given between 0 and 1. (1) *ρ*: PVE of gene expression; (2) *τ*: PVE of mediation; (3) *δ*: PVE of unmediated effect.

For each mediator *k* on individual *i*: 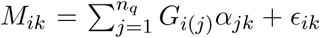 where (*j*) is *j*-th causal SNP index for this gene, *α* ~ 𝒩(0, *ρ*/*n_q_*) and ∈ ~ 𝒩(0,1 – *ρ*).

Combining stochastically generated mediator variables, we generate phenotypes

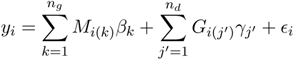

where (*k*) is *k*-th causal mediator index, mediation effect sizes are sampled independently *β* ~ 𝒩(0, *τ*/*n_g_*), and unmediated effect sizes are also sampled independently γ ~ 𝒩(0,*δ*/*n_d_*) and we infused irreducible noise to take into account of the residual variance ∈ ~ 𝒩(0,1 – *τ* – *δ*).

### Calculation of TWAS test statistics from summary z-scores

For simplicity we used completely summary-based approach^18^ that need not pretrain multivariate regression model. TWAS statistics can be derived from two sets of summary z-score vectors-one for GWAS z_*G*_ and the other for eQTL z_*T*_ using the fact that multivariate QTL effect size vector *α* can be approximated by LD-adjusted effect, 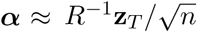.

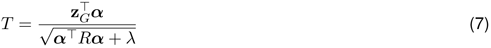

Test statistics *T* asymptotically follow 𝒩(0,1). We set λ = 10^−8^ to avoid zero divided by zero.

### Data preprocessing

We used genotypes of 672,266 SNPs in 1,709 individuals from the Religious Orders Study (ROS) and the Memory and Aging Project (MAP)^97^. We mapped hg18 coordinates of SNPs (Affymetrix GeneChip 6.0) to hg19 coordinates matching strands using publicly available information (http://www.well.ox.ac.uk/˜wrayner/strand/GenomeWideSNP_6.na32-b37.strand.zip). We imputed the genotype arrays by prephasing haplotypes based on the 1000 genome project phase I version 3^98^ using `SHAPEIT`^99^, retaining only SNPs with MAF > 0.05. We then imputed SNPs in 5MB windows using `IMPUTE2`^100^ with 100 Markov Chain Monte Carlo iterations and 10 burn-in iterations. After the full imputation, 6,516,083 SNPs were considered in follow-up expression and methylation QTL analysis.

We used gene expression data generated by RNA-seq from dorsolateral prefrontal cortex (DLPFC) of 540 individuals^4^. We retained 436 samples by first removing potentially poor quality 84 samples with RIN score below 6 (suggested by the GTEx consortium) and further removing 20 samples with no amyloid beta measurements. Of 436 samples, we used 356 samples with genotype information for eQTL analysis. Gene-level quantification was conducted by `RSEM`^101^ and we focused on 18,461 coding genes out of 55,889 according to the GENCODE annotations (v19 used in the GTEx v6p^102^).

Original gene-level quantification data follows negative binomial distribution. We adjusted variability of sequencing depth across samples following previous methods^103,104^. We then converted over-dispersed count data to Normal distribution data while fitting gene by gene null model of negative binomial regression that only includes intercept term, equivalent to `riog` transformation^105^. We used custom-designed inference algorithm to speed up the inference of models (https://github.com/ypark/fqtl). We found residual values inside the inverse link function follow approximately Normal distribution in most genes.

### GWAS summary statistics

International Genomics of Alzheimer’s Project (IGAP) is a large two-stage study based upon genome-wide association studies (GWAS) on individuals of European ancestry. In stage 1, IGAP used genotyped and imputed data on 7,055,881 single nucleotide polymorphisms (SNPs) to meta-analyse four previously-published GWAS datasets consisting of 17,008 Alzheimer’s disease cases and 37,154 controls (The European Alzheimer’s disease Initiative – EADI the Alzheimer Disease Genetics Consortium – ADGC The Cohorts for Heart and Aging Research in Genomic Epidemiology consortium – CHARGE The Genetic and Environmental Risk in AD consortium – GERAD). In stage 2, 11,632 SNPs were genotyped and tested for association in an independent set of 8,572 Alzheimer’s disease cases and 11,312 controls. Finally, a meta-analysis was performed combining results from stages 1 & 2.

## Acknowledgment

We thank Alkes Price (Harvard), Alexander Gusev (Dana Farber) and Bogdan Pasaniuc (UCLA) for insightful discussion.

We have source code for CaMMEL made publicly available as a part of the R package for summary-based QTL / GWAS analysis (https://github.com/YPARK/zqti). Effect sizes of CaMMEL are estimated through `fit.med.zqti` function.

We thank the International Genomics of Alzheimer’s Project (IGAP) for providing summary results data for these analyses. The investigators within IGAP contributed to the design and implementation of IGAP and/or provided data but did not participate in analysis or writing of this report. IGAP was made possible by the generous participation of the control subjects, the patients, and their families. The i-Select chips was funded by the French National Foundation on Alzheimer’s disease and related disorders. EADI was supported by the LABEX (laboratory of excellence program investment for the future) DISTALZ grant, Inserm, Institut Pasteur de Lille, Universite de Lille 2 and the Lille University Hospital. GERAD was supported by the Medical Research Council (Grant 503480), Alzheimer’s Research UK (Grant 503176), the Wellcome Trust (Grant 082604/2/07/Z) and German Federal Ministry of Education and Research (BMBF): Competence Network Dementia (CND) grant 01GI0102, 01GI0711, 01GI0420. CHARGE was partly supported by the NIH/NIA grant R01 AG033193 and the NIA AG081220 and AGES contract N01-AG-12100, the NHLBI grant R01 HL105756, the Icelandic Heart Association, and the Erasmus Medical Center and Erasmus University. ADGC was supported by the NIH/NIA grants: U01 AG032984, U24 AG021886, U01 AG016976, and the Alzheimer’s Association grant ADGC-10-196728.

## Author contribution

MK and PLD conceived study design. PLD provided ROS/MAP gene expression and DNA methylation data. YP, AKS and LH developed the method. YP implemented `C++` and `R` software. YP carried out mediation analysis. JD helped post-processing. YP, AKS, LH, JD, PLD and MK wrote the manuscript.

